# Dynamics of pitch perception in the auditory cortex

**DOI:** 10.1101/2024.06.10.598008

**Authors:** Ellie Bean Abrams, Alec Marantz, Isaac Krementsov, Laura Gwilliams

## Abstract

The ability to perceive pitch allows human listeners to experience music, recognize the identity and emotion conveyed by conversational partners, and make sense of their auditory environment. A pitch percept is formed by weighting different acoustic cues (e.g., signal fundamental frequency and inter-harmonic spacing) and contextual cues (expectation). How and when such cues are neurally encoded and integrated remains debated. In this study, twenty-eight participants (16 female) listened to tone sequences with different acoustic cues (pure tones, complex missing fundamental tones, and tones with an ambiguous mixture), placed in predictable and less predictable sequences, while magnetoencephalography was recorded. Decoding analyses revealed that pitch was encoded in neural responses to all three tone types, in the low-to-mid auditory cortex and sensorimotor cortex bilaterally, with right-hemisphere dominance. The pattern of activity generalized across cue-types, offset in time: pitch was neurally encoded earlier for harmonic tones (∼85ms) than pure tones (∼95ms). For ambiguous tones, pitch emerged significantly earlier in predictable contexts than unpredictable. The results suggest that a unified neural representation of pitch emerges by integrating independent pitch cues, and that context alters the dynamics of pitch generation when acoustic cues are ambiguous.

**Significance Statement:** Pitch enables humans to enjoy music, understand the emotional intent of a conversational partner, distinguish lexical items in tonal languages, and make sense of the acoustic environment. The study of pitch has lasted over a century, with conflicting accounts of how and when the brain integrates spectrotemporal information to map different sound sources onto a single and stable pitch percept. Our results answer crucial questions about the emergence of perceptual pitch in the brain: namely, that place and temporal cues to pitch seem to be accounted for by early auditory cortex, that a common representation of perceptual pitch emerges early in the right hemisphere, and that the temporal dynamics of pitch representations are modulated by expectation.

## INTRODUCTION

Pitch is central in appreciating musical melody, understanding the emotional state and intonational intent of a speaker, distinguishing lexical items in tonal languages, and correctly parsing the acoustic environment (de Cheveigné, 2010). This is enabled by the auditory system’s ability to map different sound types (e.g., a note played on a piano or a cello or sung by a human voice) onto a putatively unique and stable pitch percept.

The perception of pitch arises from a number of sources in the acoustic input. Fundamental frequency (F0), the lowest periodic component present in the acoustic signal, exhibits a strong influence on pitch perception: a pure sinusoidal tone that repeats 440 times per second will be perceived with 440 Hz pitch (de Cheveigné, 2005). Sounds arising from musical instruments or voices comprise F0 as well as a rich harmonic structure at integer multiples of F0. The spacing between harmonic components likewise exhibits a strong influence on pitch perception. Even if the energy at F0 is removed entirely, listeners perceive the signal as eliciting the same pitch percept (Seebeck, 1841; Schouten, 1938; Licklider, 1954). This “missing fundamental” phenomenon is an important clue to how the auditory system extracts pitch from auditory inputs (Plack et al., 2005; Griffiths and Hall, 2012; Wang and Walker, 2012).

How different spectrotemporal cues are integrated to solve the mapping from acoustic waveform to pitch percept, remains unknown (de Cheveigné, 2005; Oxenham, 2012). Auditory pitch determination begins at the cochlea, where frequencies are mapped to locations along the basilar membrane (Shera et al., 2002, 2010). This organization is preserved as auditory information is transmitted along the afferent auditory pathway (Oxenham and Wojtczak, 2010). The precise ways in which frequency and harmonic information are differentially represented once they reach the primary auditory cortex have been a point of contention, with tonotopy in some cases appearing to represent perceptual pitch (Pantev et al., 1989; Pantev et al., 1996; Monahan et al., 2008) and in other cases showing that the frequency of pure tones is organized tonotopically, with a separate representation of timbre (Langner et al., 1998; Warren et al., 2003; Bizley et al., 2009; Allen et al., 2017). Studies mapping pitch have typically used perceptually unambiguous, narrowband signals as pitch-evoking stimuli (pure tones, ripple-induced noise, relative intensity noise), making it unclear whether these representations track perceptual pitch invariant to spectral structure (Allen et al., 2022) or whether they reflect an F0 extraction mechanism (Kumar et al., 2011b; Griffiths and Hall, 2012). To further complicate this issue, populations of neurons in immediate proximity to spectrally invariant pitch-selective neurons have been shown not to respond to pure tones, but to respond to complex tones and other narrow- or wideband stimuli (Bendor and Wang, 2005). Regarding spectral structure, some auditory neurons are sensitive to spectral contrast, or the difference in intensity between frequency components in wideband stimuli, highlighting spectral shape as a crucial dimension across which stimuli may be represented (Barbour and Wang, 2003). Lastly, listening context (i.e., interspersing stimuli with silence vs. noise) may modulate pitch sensitivity for some auditory regions, but not others (Garcia et al., 2010).

Pitch-selective cortical regions, or, “pitch centers,” have been established in lateral Heschl’s gyrus, planum temporale, and superior temporal gyrus (Patterson et al., 2002; Hall and Plack, 2009; Gander et al., 2019; Kim et al., 2022), as well as in anterior non-primary areas (Penagos et al., 2004). This diversity of cortical representation could be driven by stimulus properties, with some areas demonstrating a preference for resolved harmonics (Norman-Haignere et al., 2013) or reflecting one specific stimulus feature (Barker et al., 2011). Alternatively, these areas could comprise a distributed system of neural ensembles representing pitch over auditory cortex (Griffiths et al., 2010; Kumar et al., 2011a; Allen et al., 2022). The use of more temporally resolved methods (EEG/MEG/local field potentials) has allowed for an exploration of pitch perception at the millisecond level, but work has focused on singular evoked responses as opposed to the unfolding of a percept over time (Brattico et al., 2001; Loui et al., 2009; Butler and Trainor, 2012; Kim et al., 2022).

An additional cue to pitch perception is the context in which a sound occurs (Krumhansl and Shepard, 1979; Krumhansl and Cuddy, 2010; Slana et al., 2016), which has been shown to bias perception towards a contextually appropriate interpretation when two acoustic cues are ambiguous or in conflict (Shepard, 1964; Deutsch, 1992; Deutsch et al., 2008). Further, when there is a high tonal expectation of the next pitch, the expected pitch is neurally represented even when omitted from the end of a sequence (Berlot et al., 2018). This effect is demonstrably present as early as primary auditory cortex (Englitz et al., 2024), suggesting that context is a fundamental component of cortical pitch encoding. However, it is unknown how contextual expectation interacts with acoustic content to derive an ultimate pitch representation.

In the current study, we aim to disentangle these three primary cues to pitch perception: F0, inter-harmonic spacing, and tonal context. Magnetoencephalography (MEG) data were collected from 28 healthy adults presented with sequences of pitch-matched tones with varying spectral content in and out of context. Pure tones and missing fundamental (MF) complex tones were used to represent F0 pitch cues and inter-harmonic spacing pitch cues, respectively. Ambiguous tones were used to investigate the neural representation of pitch in acoustic ambiguity. Decoding analyses were applied to the MEG data to test hypotheses about shared representations between tone types. The results provide new evidence for the cortical timing associated with the analysis of these basic cues to pitch.

## MATERIALS AND METHODS

### Participants

We recruited 28 participants from the NYU Abu Dhabi community (16 females, mean age 24.8 years). All subjects reported normal hearing and no history of neurological disorders. Participants were compensated for their time at a rate of 60 dirhams per hour. The experiment was approved by the NYU Abu Dhabi IRB committee.

### Stimuli

Pure tones consisted of one frequency band at F0 with no additional harmonic components, for a total of 17 unique frequencies (Figure 1A, D). Complex missing fundamental (MF) tones consisted of the five integer multiples (partials) above, but not including the fundamental frequency for each of the 18 pitches (Figure 1A, D). For example, a complex MF tone with energy at frequencies 440, 660, 880, 1100 and 1320Hz has a distance of 220Hz between each harmonic and will be perceived as having a pitch of 220Hz; yet the 220Hz frequency content itself is not present. These missing fundamental tones had a flat spectral envelope, with equal attenuation of all five harmonics.

**Figure 1.**
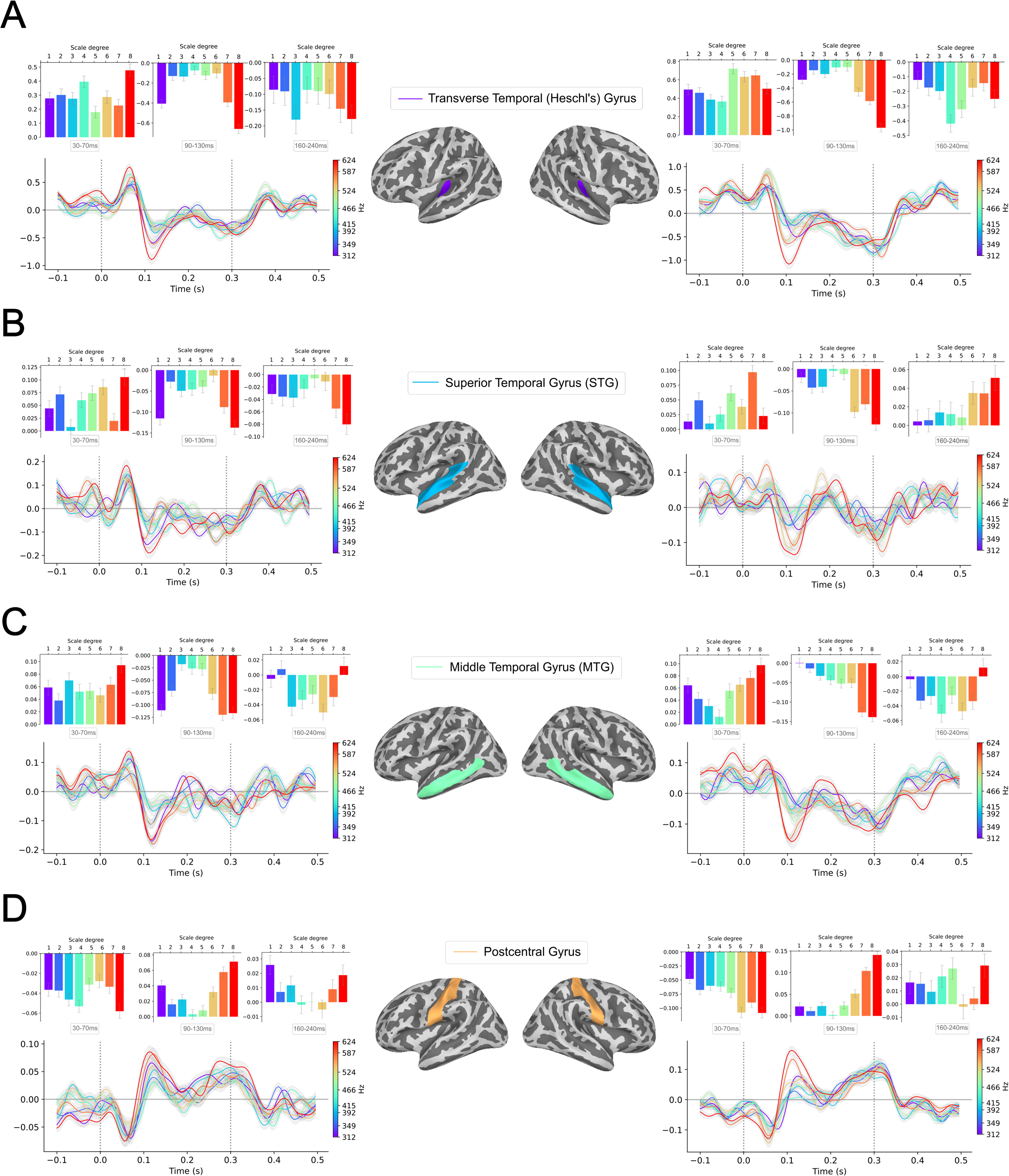
Experimental stimuli. **(A) Tone types.** Waveforms, spectral content, and perceived pitch for a pure tone, complex missing fundamental (MF) tone, and ambiguous tone of 220Hz. **(B) Evoked responses.** MEG evoked responses (and topoplots) across all sensors averaged across participants, for each tone type. **(C) Musical keys.** Musical notation for each of the three scales used to create stimuli. **(D) Harmonic components.** Harmonic components for each scale degree of one example scale (A major) for each stimuli type. Pure and complex MF tones contain flat amplitudes across harmonics, while ambiguous tones contain differently attenuated harmonics for each scale degree.

To build ambiguous tones, we use Diana Deutsch’s adaptation of the Shepard tone illusion, in which she systematically varied the amplitudes of odd and even harmonics across the semitones spanning one musical octave, resulting in a set of 7 tones corresponding to the notes of a diatonic scale. Due to the ambiguity of the tones, when they are played in succession repeatedly, the sequence of tones evokes an auditory barbershop effect, sounding like a continuously ascending or descending scale with no stable perceived switch during the point at which the tones repeat. Because of competing pitch cues, namely the actual fundamental frequency in tension with the spacing between harmonics, the tonal center (i.e., the key of the scale) may be perceived as either the low or high octave of the scale. Concretely, as the tones decrease in fundamental frequency, the even harmonics remain the same in amplitude, while odd harmonics decrease. This leads to competing cues to pitch, namely, a ratio of 2*F0 between harmonics and a highly attenuated, albeit present, F0. Though participants were only presented with the 7 semitones making up the major scale, 12 complex MF tones were created to maintain equal stepwise attenuation of the odd harmonics. Each contained the first six partials (multiples of F0). The parameters used to achieve ambiguity, starting at the top of the scale, were: Partials 2 and 4 remained flat for all tones, while partials 1 and 3 were attenuated by 3.5 dB with each semitone decrease in f0. Partial 5 was attenuated to 28 dB for all semitones. Lastly, partial 6 began at 28 dB and was strengthened by 3.5 dB with each semitone decrease in F0 (Figure 4D).

Pure, missing fundamental and ambiguous tones were synthesized in three musical keys: A3 (220-440Hz), C4 (262-524Hz), and Eb4 major (312-624Hz; Figure 1C). Frequency values were calculated according to the equal tempered scale, in which semitones are separated by constant frequency multiples. These frequencies ranged from 220-624 Hz. These three banks of tones, corresponding to the three experimental blocks (pure, complex MF, and ambiguous), were created using PyAudio, a python package for playing and recording audio on Mac OS X. Tones were saved as wav files with a sampling rate of 44,100 Hz. All tones were 300ms in duration, 32 bits per sample, and normalized to 70 dB using Praat.

## EXPERIMENTAL DESIGN AND STATISTICAL ANALYSES

All tones were presented for a duration of 300ms, with a 200ms silent inter-stimulus interval. Pure and complex MF blocks each contained nine mini-blocks: up, down, and random for each of the three musical scales. Each mini-block repeated the 8 tones 15 times, with tone order depending on the condition. In the up and down conditions, the tones were presented in ascending and descending order, respectively, with a jittered 500-750ms of silence in between each mini-block. In the random condition, the 8 tones were presented in random order 15 times. All mini-blocks were shuffled within larger pure tone and complex MF tone blocks.

Participants were presented with the ambiguous tones in a scale condition which mirrored that of the pure and complex MF tones. Because there are only 7 ambiguous tones per scale, we played the ambiguous tone at both the beginning and end of the scale so that it would explicitly act as the low and high octave (Figure 1D). The scale condition was presented in ascending (up condition; Figure 1C) and descending (down condition) order. As in the pure and complex MF scale conditions, repetitions were separated by a pause of 500-750ms. Lastly, during the random condition, the 7 tones were presented in random order 15 times. These 15 mini-blocks were shuffled.

All tones were presented binaurally to participants through earphones in the magnetically shielded room (MSR). Participants listened passively, while watching silent videos of nature scenes. Eye tracking was used to ensure participants were not falling asleep. Block order was counterbalanced across participants. The experiment lasted around 80 minutes.

### Data acquisition and preprocessing

Prior to recording, each subject’s head shape was digitized using a Polhemus dual source handheld FastSCAN laser scanner. Digital fiducial points were recorded at five points on the individual’s head: the nasion, anterior of the left and right auditory canal, and three points on the forehead. Marker coils were placed at these same points to localize each subject’s skull relative to the MEG sensors. To correct for movement during recording, measurements from the marker coils were obtained immediately before and after the protocol.

Continuous MEG data were recorded using a 208-channel axial gradiometer system (Kanazawa Institute of Technology, Kanazawa, Japan) with an online bandpass filter of 0.1-200Hz at a sampling rate of 1000Hz. Continuous recording was noise reduced using Continuously Adjusted Least Squares Method (CALM) (Adachi et al., 2001), with MEG160 software (Yokohawa Electric Corporation and Eagle Technology Corporation, Tokyo, Japan). The noise-reduced MEG recording, digitized head-shape, and sensor locations were then imported into the Python package *mne* (Gramfort et al., 2013). After visual inspection, one saturated channel was rejected in two subjects. No channels were rejected in the remaining subjects after visual inspection. An independent component analysis was fitted on the data to account for 90% of explained variance, and components corresponding to EKG, eye movements, and blinks were removed.

Epochs were extracted from −200ms pre-stimulus onset to 600ms post-stimulus onset, and down-sampled to 200 Hz. Individual epochs were rejected using an absolute threshold of ±2000 femto-tesla; less than 5% of epochs were rejected in all subjects. See Figure 1B for evoked responses averaged across subjects, for each experimental condition.

### Source Localization

To perform source localization, the location of the subject’s head was co-registered with respect to the sensory array in the MEG helmet. Structural Magnetic Resonance volumes (T1s) were collected from all participants using NYUAD’s MRI facility (3T MAGNETOM Prisma, Siemens). These anatomical scans were rotated and translated using *mne* to minimize the distance between the fiducial points of the MRI and the head scan.

Next, a source space was created, consisting of 2562 potential electrical sources per hemisphere using an “ico-4” source space. At each source, activity was computed for the forward solution with the Boundary Element Model (BEM) method, which provides an estimate of each MEG sensor’s magnetic field in response to a current dipole at that source. The inverse solution was computed from the forward solution and the grand average activity across all trials. Data were converted into noise-normalized Dynamic Statistical Parameter Map (dSPM) units (Dale et al., 2000), using an SNR value of 2. The inverse solution was applied to each trial at every source, for each millisecond defined in the epoch, employing a fixed orientation of the dipole current that estimates the source normal to the cortical surface and retains dipole orientation.

### Multivariate decoding analysis

We implemented our decoding analysis using the Python packages *mne* (Gramfort et al., 2013) and *scikit-learn* (Pedregosa et al., 2011).

We regularize model fit using L2 regularization, with a fixed strength of alpha=1e+8.

Before fitting the decoder, we normalize the magnitude of the MEG sensors using *StandardScaler*, a function of *scikit-learn*:

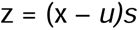

where *u* is the mean of the training samples and *s* is the standard deviation of the training samples.

For cross validation, we first randomly shuffle the order of trials. Then we subset the data into five partitions of 80% - 20% splits, with non-overlapping trials across split assignment. This was implemented using the KFold (k=5) cross validation scheme in *scikit-learn*. The 80% of trials we used to train the decoder (i.e., learn the coefficient weights that optimally map the MEG sensor activity to the stimulus feature), and the remaining 20% of the trials to generate stimulus predictions based on the learned coefficient maps. We repeat this fit-predict procedure for each of the 5 folds of the data and average the decoding performance across the five folds.

Decoding analyses were applied to the 200 Hz bandpass-filtered epochs. The shape of the neural data matrix per subject was number of trials (7650) x number of MEG sensors (208) x number of timesamples from −200:600ms at 200 Hz (161) x number of subjects (28). This neural data matrix was used for all decoding analyses.

We also derive a corresponding stimulus feature vector, which contains the same number as trials as the neural data. For example, the stimulus vector that defines musical pitch includes all the pitch values of each trial (e.g., 440Hz, 312 Hz, etc.); the stimulus vector that defines categorical tone type includes the tone type of each trial, such as “pure tone”, “complex MF tone”, etc.

To address the question of whether a stimulus feature of interest is present in neural responses, we derived a *decoding accuracy timecourse*. For categorical stimulus features, such as tone-type, we use a Logistic Regression classifier. In this case, we transform the category labels into binary [0, 1] sequences, and use one-versus-all Logistic Regression. The performance of this classifier is quantified using receiving operator characteristic area under the curve (ROC AUC), by comparing the true binary property of the feature [0, 1] to the probabilistic output of the classifier, bounded between 0-1. Chance-level performance of AUC correlation is 0.5.

For continuous stimulus features, such as pitch, we used Ridge Regression. We scaled the stimulus values linearly between 0-1, such that the lowest value is assigned 0, the highest value is assigned 1, and the relative distance between values is preserved in the scaled space. We computed performance using a Spearman rank correlation, which compares the continuous true property of the feature to the continuous predicted property of the feature. Chance-level performance of a Spearman correlation is 0. Because a rank correlation metric was used, the distances between frequencies did not matter. Thus, using log scale as opposed to linear would not affect decoding performance.

Decoding models were fit for each subject independently, and a separate model was fit at each time-lag relative to tone onset. That means that we fit a total of 161 x 28 = 4,508 models. The input to each model is the strength of neural activity in units of femto-tesla across all MEG sensors across trials.

The step-by-step procedure to derive a decoding timecourse for each subject is as follows: we loop through each timepoint (161) and K fold (5 subsets of training and testing epochs) and fit a decoder on the MEG activity (7650 trials total; X) and corresponding stimulus features (tone-type or pitch; y) at each *training epoch* at that timepoint. We use this classifier to predict stimulus features for unseen *testing epochs* at that same timepoint. Lastly, we compute performance by comparing the prediction output by the decoder with the true feature of the *testing epochs*.

To address the question of how information is spatially encoded in the brain, we can similarly derive a *spatial* map of decoding performance by using the same analysis logic, but transposing space and time. In other words, whereas to derive a decoding timecourse we fit a decoding model at each time-step separately and used the activity over sensors as the neural features, we can instead model each reconstructed neural sources separately and use the strength of activity across time-steps as the neural features. We decided to focus this analysis on the 100ms around the peak of acoustic decoding (see Source Localization, above).

The step-by-step procedure for spatial decoding is like that of sensor space decoding, but instead of looping through timepoints we loop through each spatial source and fit the decoder on activity during 50ms windows throughout the entirety of the epoch (−200 - −150ms through 550-600ms).

### Second-order statistical tests

To assess statistical significance of decoding analyses in sensor space, we compared the distribution of decoding performance across subjects against chance performance. Concretely, we formulated a matrix of shape nsubjects (28) x ntimes (161) and fit a one-sample permutation cluster test to derive putative clusters of significance (implemented in *mne.stats.permutation_cluster_1samp_test* in *mne-python).* This test first performs a t-test for the distribution of decoding performance across subjects compared to chance level (0.5 for AUC and 0 for Spearman). This array of t-values is then clustered based on adjacent values (either in time, or in space, depending on the analysis), that exceed a significance threshold of t > 1.96 (equivalent to p < .05). Then, the average t-value in the cluster is recorded. Finally, surrogate data are formed by randomly flipping the sign of the input data. So, the score will be randomly assigned a positive or negative value, arbitrarily above or below chance. Then, the t-tests and clusters are re-formed based on this chance-derived surrogate data. This surrogate procedure is repeated 1000 times to form a distribution of average t-values for clusters. We then count the number of times that a surrogate t-value magnitude was equal to or greater than the critical observed t-value in the data. This count divided by the 1000 surrogate values forms our statistical p-value. For example, to derive a p-value of p < .05, there would be no more than 50 surrogate t-values that exceeded the empirical data.

To evaluate statistical significance of source decoding within an ROI, we extract the R/AUC values within each ROI for each subject, for each hemisphere. The statistical significance of decoding in a given ROI is calculated by averaging the R/AUC values across sources in the ROI, and then performing a one-sample t-test of the Spearman R or AUC values of these sources against chance (0 or 0.5, respectively), and a paired t-test comparing scores between hemispheres.

## RESULTS

Our use of stimuli with non-overlapping spectral content allowed us to take advantage of the ability for F0 (*pure tones)*, on the one hand, and inter-harmonic spacing F0 (*complex missing fundamental (MF) tones*), on the other, to convey pitch-relevant information *independently*. Our use of stimuli with an ambiguous mixture of acoustic cues (*ambiguous tones*), which elicit distinct pitch percepts depending on context (Deutsch et al., 2008), allowed us to test the effect of unambiguous versus ambiguous acoustic inputs and the effect of predictable and unpredictable contexts, on the neural encoding of pitch. Importantly, although the missing fundamental has been shown to be restored perceptually, and potentially represented weakly in brainstem auditory nuclei, at the level of the cochlea the two stimulus types are non-overlapping: F0 is represented along the basilar membrane in pure tones, but not in MF tones, even as a non-linear distortion product (Greenberg, 1980; Galbraith, 1994). However, the extent to which these differing pitch cues have been processed or represented for perceptual interpretation at the level of cortex is unclear. Using a decoding framework, we address three questions: First, is pitch encoded differently for sounds with different spectral content (reflecting the input), or is there a shared pitch representation (reflecting the percept)? Second, if there is a shared neural representation, when and where is that representation generated? Third, when acoustic pitch cues are ambiguous, how is the pitch representation affected by contextual expectations?

### Evoked ROI activity

Tones were presented in blocks corresponding to three different musical keys, in randomly presented order or in scale order. The scale order condition is useful in ensuring a contextual expectation for the next tone, but evoked representations of the low and high octave of the scale may be difficult to interpret due to their position at the beginning or end of the scale sequence. Thus, in Figure 2, we show evoked activity in four regions of interest (ROIs; Heschl’s gyrus, superior temporal gyrus, middle temporal gyrus, and postcentral gyrus) for the 8 scale degrees of one of the musical scales (Eb Major), presented randomly. The three auditory ROIs were selected *a priori*, as motivated by prior literature investigating pitch processing in auditory cortex (Patterson et al., 2002; Hall and Plack, 2008; Wilson et al., 2009). The somatosensory ROI was a data-driven inclusion, based on our observation of strong tone-evoked activity in this region. We extract the ROI activity using the *aparc* parcellation.

**Figure 2.**
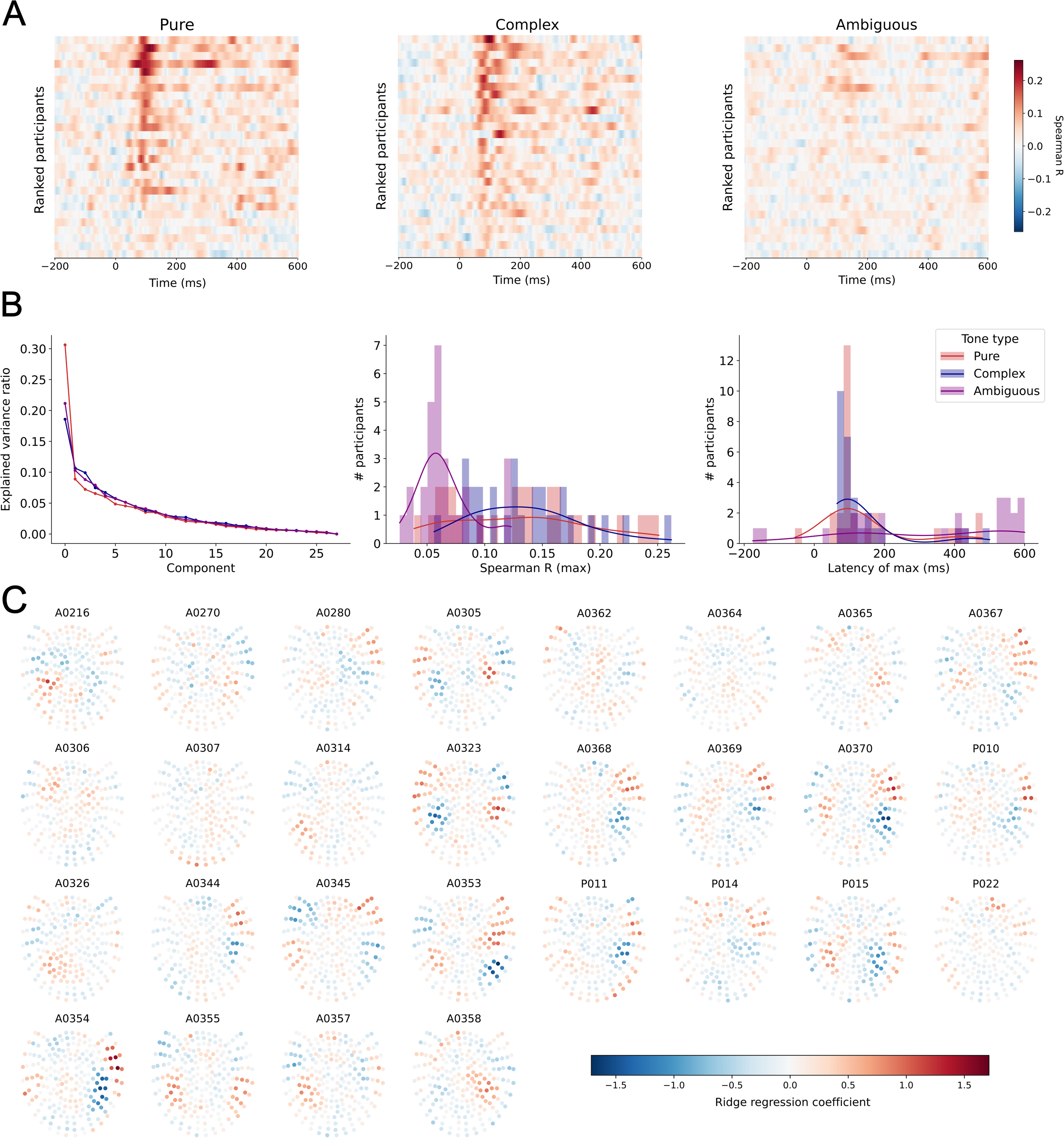
Evoked ROI activity. MEG activity averaged across all trials and all participants for the 8 scale degrees of the Eb Major scale, tones presented randomly. Activity is plotted in the left (*left column)* and right *(right column)* hemisphere for four ROIs: transverse temporal (Heschl’s) gyrus **(A)**, superior temporal gyrus **(B)**, middle temporal gyrus **(C)**, and postcentral gyrus **(D)**. Bar plots show average activity within three temporal windows after tone presentation: 30-70ms *(left bar plot)*, 90-130ms *(middle bar plot)*, and 160-240ms *(right bar plot)*.

During the window around M100-associated activity, average activation in right hemisphere ROIs (Figure 2, right column) seems to increase with scale degree, which indicates a tonotopic, F0-based organization in terms of amplitude. However, the left hemisphere shows comparable amplitudes between the first and last scale degree (low Eb and high Eb; Figure 2, left column), suggesting an effect of pitch chroma, or octave equivalence. This is in accordance with work showing distinct mappings of pitch height (specific octave) vs. chroma (cyclical notes of a scale) (Warren et al., 2003) and an underlying octave structure to tonotopy (Brosch et al., 1999; Brosch and Schreiner, 2000). However, this left lateralization is novel, as octave-tuned voxel clusters have been found mostly in the right hemisphere (Moerel et al., 2015), and bilateral chroma sensitivity has been shown for complex, but not pure tones (Briley et al., 2012). The postcentral gyrus appears to represent pitch in a similar way to auditory areas, which will be discussed later.

### Individual decoding differences and reliability

Before aggregating individual responses in our decoding analyses, we first look at individual pitch decoding timecourses (Figure 3A). A general pattern emerged of performance peaking within the first 100ms for pure and complex tones. The peak is much less pronounced for ambiguous tones, but the pattern is consistent across subjects. There are, however, some subjects that exhibit notable double peaks and others that do not seem to peak at all.

**Figure 3.**
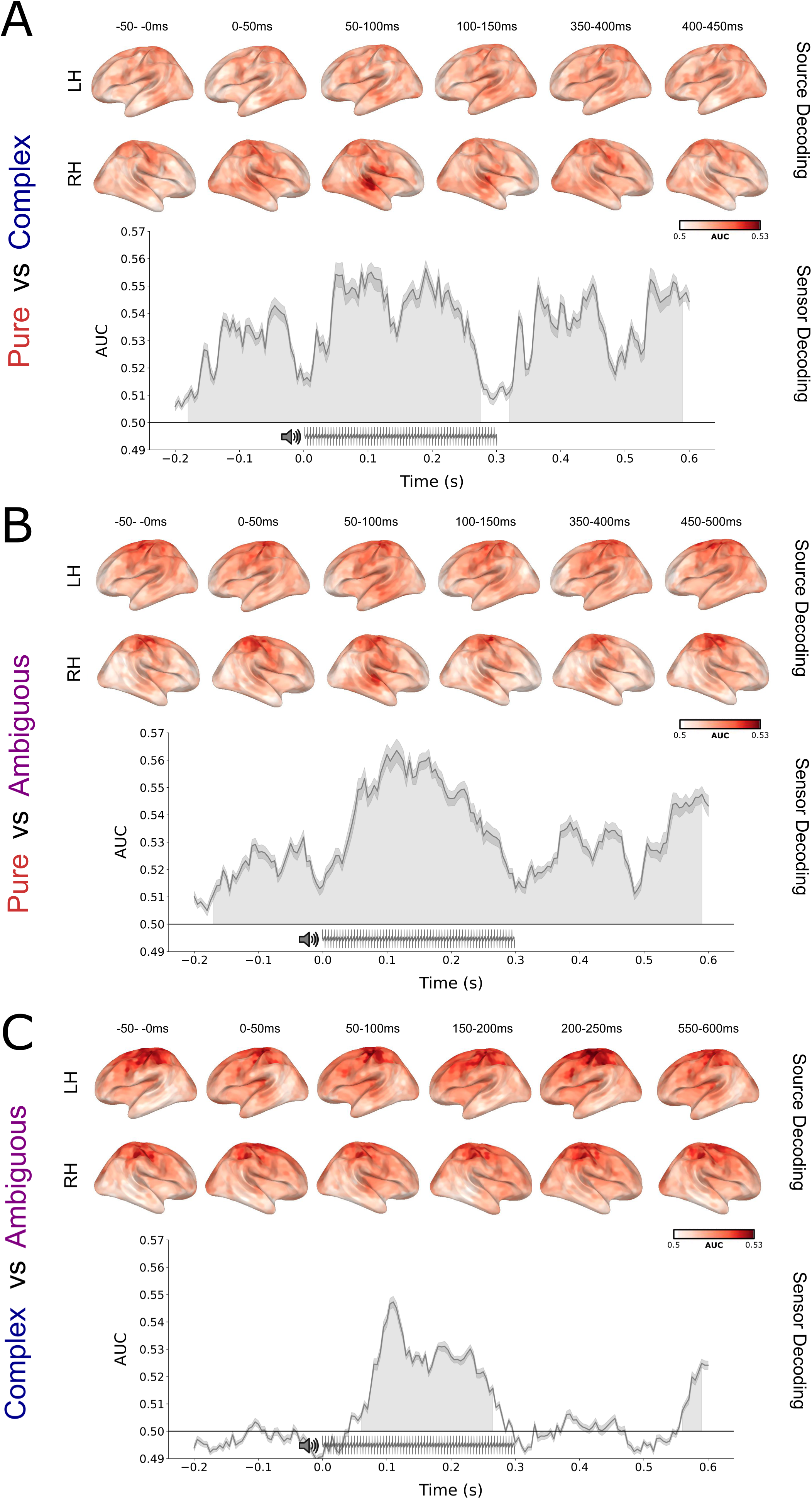
Individual differences and reliability of decoding. **(A)** Individual decoding performance for pure tones *(left),* complex MF tones *(middle),* and ambiguous tones *(right).* Each row represents one participant, which are ordered by maximum decoding performance across the three tone types. **(B)** Principal components for each decoding timecourse *(left)*; Distribution of maximum decoding scores, plotted as a histogram with bin width 0.006, for each tone type. This is overlaid with a Gaussian kernel density estimator (KDE) approximation with bandwidth selected via Scott’s method (Scott, 1992) *(middle)*. Latency of max Spearman R, plotted with a bin width of 20, overlaid with a Gaussian KDE to select bandwidth *(right)*. **(C)** Feature importance sensor maps for all participants for pure tone decoding, plotted as Ridge regression weights for each sensor at decoding peaks averaged across five folds of cross-validation.

We next performed a principal components analysis (PCA) on these timecourses and plotted the proportion of explained variance for the first 28 components (Figure 3B, left). For all tone types, there was a pronounced 1-factor structure, with explained variance leveling off beyond the first component. This means that much of the variance in individual responses could be explained by the scaling of a single response profile. In addition, we measured the distribution of maximum pitch decoding scores (Spearman correlations) for each tone type within the first 200ms following tone onset (Figure 3B, middle), as well as the time of each maximum score, relative to tone onset (Figure 3B, right). Together, these show that although subjects varied in their maximum decoding scores, the latencies of these maxima are highly consistent across tone types.

Finally, we examined which MEG sensors contributed most to pitch decoding in the pure tones condition for each subject (Figure 3C). We did this by extracting the Ridge regression weights assigned to each sensor when decoding performance peaked, averaged across five folds of cross-validation. The weights vary topographically by subject, but most show clusters of high-contributing sensors in bilateral temporal regions, with the upper and lower halves of each region showing opposite polarity. Taken together, these analyses suggest that there is variability in the strength, but not the structure, of pitch encoding as derived from individual MEG activity.

### Decoding tone type

Next, we tested at the group level whether pure and complex MF tones elicit distinct auditory responses, given that their spectral content is different, but their fundamental frequency is the same. To do so, we used a logistic regression classifier to distinguish between the pure and complex MF ‘tone types’ (see Methods). We fit the classifier at each 5-millisecond time sample separately, for each subject, using the activity across the 208 MEG channels for each epoch as the input features. This resulted in a timecourse of decoding performance for each subject, resolved at the 5-millisecond scale. Classifier performance was quantified using area under the curve, which summarizes the similarity between the true stimulus labels and the classifier’s predictions of those labels. If the two signal types elicit the same neural response, the classifier will be unable to provide an accurate label to distinguish the two classes, and so the decoding performance should not differ from chance (AUC=0.5). If the two tones systematically elicit two distinct responses, the difference in response will be learnable by the classifier, leading to above-chance decoding performance. To evaluate statistical significance, we used a temporal permutation cluster test (see Methods). This nonparametric approach controls for Type I error by empirically testing the distribution of results that could have been obtained by chance and comparing that to the observed result. The output is a series of time-windows that have been evaluated for statistical significance. Given that the tone was played from 0 to 300ms, we used a search window of −200 to 600ms, to investigate neural responses during the stimulus and during the silence in between tones.

As shown in Figure 4A (left panel), tone type for pure versus complex MF tones could be decoded significantly better than chance for almost the entire duration of the tone (−180-280ms, df = 27; t = 5.2, p = 0.0001) as well as the silent period before the next tone (320-525ms, df = 27; t = 4.98, p = 0.0001). This suggests that spectral distinctions between acoustic inputs are robustly encoded throughout the processing of the tone and are even maintained between tones.

**Figure 4.**
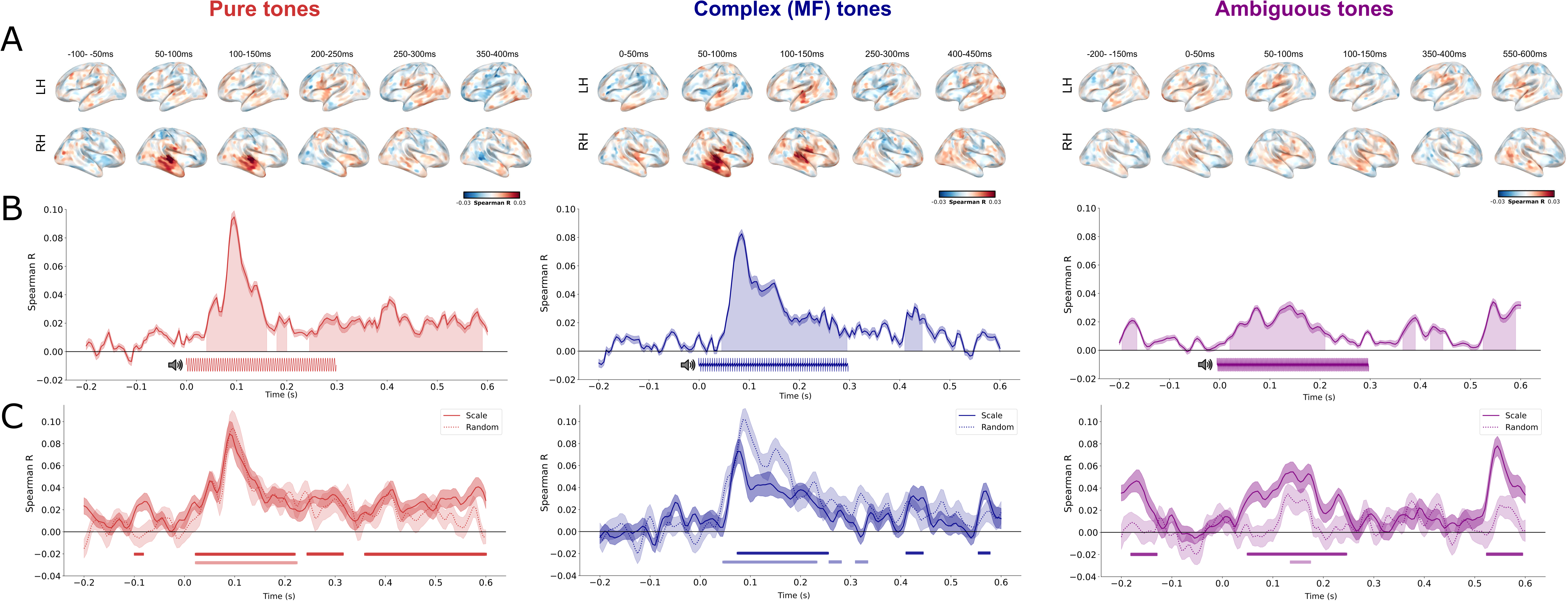
Tone type decoding. Decoding performance over time is plotted with standard error of the mean for each combination of tone types: pure vs. complex MF (*top*), pure vs. ambiguous (*middle*), and complex MF vs. ambiguous *(bottom*). Temporal clusters of significant above-chance decoding (AUC) are shown shaded under the curve. Stimulus waveforms are depicted below each timecourse to denote tone duration. Source decoding of tone type averaged across 50-150ms after tone onset is shown for each tone type comparison. Colorbar denotes AUC values for each source.

To determine the localization of the missing fundamental (MF) spectral distinction, we first performed source localization on the MEG single trial epochs. This transformed the data from a 208-dimensional sensor space into a 5124-dimensional source space, with reconstructed sources tiling the cortical surface. We then morphed the activity projected onto each subject’s MRI to a common template brain, to have the same source space across subjects. To determine which of these localized sources encoded the spectral distinction between tone types, we again used a decoding framework. Instead of fitting the logistic regression classifier with the activity pattern across space at each millisecond, we instead fit the classifier with a window timecourse of activity at each reconstructed source. This yielded a spatial map of AUC scores, for each spatial source, for each subject.

We fit this decoder on the timecourse of activity in 50ms windows for the entirety of the epoch to produce a spatial map of decoding performance across time (Figure 4). We found that the most informative sources for distinguishing pure and complex MF tones throughout the entirety of the tone and silent periods (Figure 4A) were in the auditory cortex, bilaterally within Heschl’s gyrus (df = 27, average t = 5.31, p < 0.001), superior temporal gyrus (df = 27, t = 5.4, p < 0.001) and middle temporal gyrus (df = 27, average t = 4.27, p < 0.001), and in the sensorimotor cortex, within the precentral gyrus (df = 27, average t = 3.19, p < 0.01) and postcentral gyrus (df = 27, average t = 3.42, p < 0.001). Right hemisphere sources were more significant than left hemisphere in superior temporal gyrus (df = 27; average t = 2.28, all p < 0.05) and postcentral gyrus (df = 27; average t = 2.37, all p < 0.05) from 0-100ms and in Heschl’s gyrus from 0-50ms (df = 27; t = −3.15, p = 0.004). Tone type encoding in the postcentral gyrus was also right hemisphere dominant between 250-350ms (df = 27; average t = 2.64, all p < 0.03). There was no significant difference in tone type encoding between auditory and sensorimotor areas. This suggests that pitch-invariant spectral envelope structure is encoded in neural populations throughout the auditory and sensorimotor cortex, with a right-hemisphere dominance.

Finally, we sought to understand to what extent the neural responses to pure and complex MF tones were distinguishable from the ambiguous tones, given that we created them as a weighted mixture of acoustic cues. We fit the same logistic regression classifier described above. Decoding performance for pure vs. ambiguous tones was similar to that of pure vs. complex MF, with tone type being decoded significantly better than chance for the entire duration of the tone, including the silent period after the tone (−170-590ms, df = 27; t = 4.80; p = 0.0001) (Figure 4B). Next, we trained a classifier to distinguish between complex MF and ambiguous tones. Here, we found that tone type was decoded significantly better than chance only for the duration of the tone (60-270ms, df = 27; t = 6.78, p = 0.0001), as well as a short period directly before the beginning of the next tone (560-595ms, df = 27; t = 3.45, p = 0.01; Figure 4C). This attenuated distinction between complex MF and ambiguous tones makes sense, given that their spectral complexity and composition is similar compared to the pure tones. Indeed, the ability to distinguish them at all is quite remarkable given the subtlety of their acoustic difference.

We evaluated the spatial localization of these contrasts using the same decoding procedure we applied for pure and complex MF tones. Interestingly, the localization of effects was similar, but with sensorimotor area dominance. The distinction between pure and ambiguous tones localized to auditory cortex and sensorimotor cortex. Heschl’s gyrus encoded this distinction only in the left hemisphere from −50-0ms (df = 27; t = 3.88, p = 0.0007) and from 150-300ms (df = 27, average t = 3.24, all p < 0.01), but bilaterally otherwise (df = 27; average t = 4.14, all p < 0.01). Tone type was significantly encoded in bilateral superior temporal gyrus (df = 27, average t = 3.81, all p < 0.01), middle temporal gyrus (df = 27, average t = 3.52, all p < 0.01), precentral gyrus (df = 27, average t = 3.51, all p < 0.01) and postcentral gyrus (df = 27, average t = 3.81, all p < 0.001). There was significantly higher decoding in sensorimotor areas than auditory areas from −200-50ms (df = 27, average t = 3.5, all p < 0.01) and from 450-550ms (df. =27; average t = 3.44, all p < 0.001).

The more subtle distinction between complex MF and ambiguous mixture tones localized to bilateral sensorimotor cortex for the entirety of the tone and silent periods, with significant decoding in precentral gyrus (df = 27, t = 2.98, p = 0.01) and postcentral gyrus (df = 27, t = 3.50, p < 0.001). Tone type decoding in auditory areas was left hemisphere dominant during early and later windows (−200-0ms, 350-600ms; df = 27; average t = 2.45, all p < 0.01) but bilateral middle and superior temporal gyri encoded pitch from 0-350ms (df = 27; average t = 3.45, all p < 0.0005). Heschl’s gyrus encoding was significantly higher in the left hemisphere from 50-100ms (df = 27; t = 2.35, p = 0.03) and left hemisphere dominant from 100-500ms (df = 27; average t = 3.10, all p < 0.01). Decoding was more significant in sensorimotor areas than auditory areas for all time windows except for 50-100ms (df = 27, average t = 2.47, all p < 0.03).

Although representing distinct information, the pre- and postcentral gyri are functionally integrated, with regions of high connectivity linking somato-cognitive control and action (Gordon et al., 2023). Our initial findings support the notion that pre- and postcentral gyri together may comprise an important node of our auditory processing apparatus. The structure of responses in motor cortex have been shown to represent acoustic features over articulatory features when listening to speech sounds (Cheung et al., 2016), and melody and tempo discrimination tasks have been shown to engage the postcentral gyrus specifically (Thaut et al., 2014). Additionally, although often reflecting task-related motor processing, sound-related motor representations have been found in SMA and pre-SMA during passive listening tasks and may represent sensory predictive processing during auditory perception (Zatorre et al., 2007; Nastase et al., 2014; Lima et al., 2016; Gordon et al., 2018). Differences between musicians and non-musicians, as well as between those with and without absolute pitch, have been shown in premotor areas, with musicians recruiting pre-SMA and SMA more than non-musicians during music perception tasks (Grahn and Rowe, 2009; Margulis et al., 2009). Specifically relevant to the distinction between ambiguous tones and pure/complex tones, sensorimotor cortex has been implicated in the processing of timbre (Wallmark et al., 2018; Criscuolo et al., 2022). Taken together, the initial pitch organization we find in evoked responses (Figure 2) as well as the encoding of subtle spectral differences (Figure 4) are supported by these results.

### Decoding pitch from distinct acoustic cues

Having established that the responses to the three tone types are distinct, we next focused on the encoding of pitch in each of the three tone types. We again used a decoding framework for this. Because pitch is a continuous feature rather than a binary class, we used a ridge regression model. Accordingly, we used Spearman rank correlation to summarize the similarity between the true pitch values and predicted pitch values, given that they are both continuously varying. Chance performance for a Spearman correlation is zero.

We performed the decoding analysis separately for each tone type and derived a timecourse of Spearman correlation values for each subject. We then performed statistical analyses on the distribution of decoding performance across subjects to test whether it was significantly different from chance. Pitch decoding was significant for pure tones between 40 –165ms (df = 27; t = 5.67, p = 0.0001) and during the silent period between 245-595ms (df = 27; t = 3.25, p = 0.0001; Figure 5B, left panel). This final period includes stimulus-related evoked activity, as well as the silent period following each tone. For complex MF tones, pitch decoding was significant from 45-300ms (df = 27; t = 4.8; p = 0.01) and from 410-450ms (df = 27; t = 4.07; p = 0.008; Figure 5B, middle panel). Decoding performance peaked at 95ms for pure tones, and at 85ms for complex MF tones. Overall, this result demonstrates that pitch is robustly encoded in neural responses to the two spectrally non-overlapping tones.

**Figure 5.**
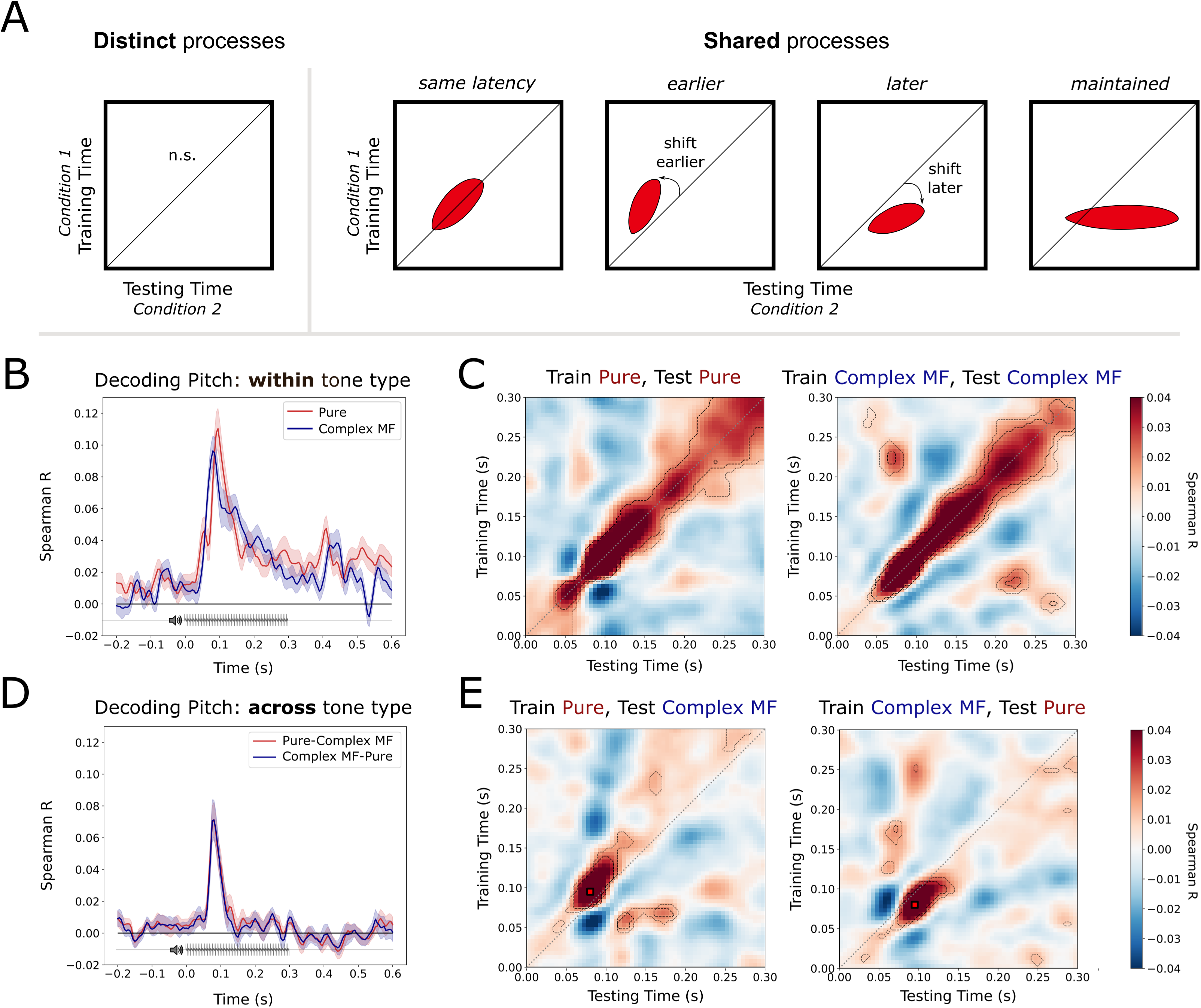
Pitch decoding. **(A)** Source space decoding of pitch averaged across selected 50ms windows. Colorbar denotes Spearman R values for each source. **(B)** Decoding performance over time plotted with standard error of the mean for each tone type: pure (*left*), complex MF (*middle*), and ambiguous (*right*). Temporal clusters of significant above-chance decoding (Spearman correlation) are shown shaded under the curve. Stimulus waveforms are depicted below each timecourse to denote tone duration. **(C)** Decoding performance for pitch in tones within context (scale, solid line) and out of context (random, dotted line) for each tone type. Shaded areas show standard error of the mean (SEM). Bars below show significant above-chance decoding clusters (Spearman correlation).

Having established that two non-overlapping acoustic cues for pitch – F0 and inter-harmonic spacing – lead to robust neural encoding of pitch along a similar timecourse, we next tested for the encoding of pitch in the ambiguous tones. Although pitch is ambiguous, leading at times to distinct percepts depending on context, we assigned pitch labels based on the lowest present frequency, F0. Pitch was significantly decodable from −195 – −160ms (df = 27; t = 4.04, p = 0.02), 20-215ms (df = 27; t = 3.9, p = 0.0001) with notable a dip in decoding accuracy at 95ms, 365-395ms (df = 27; t = 2.9, p = 0.05), and 525 – 595ms (df = 27; t = 4.05, p = 0.0007; Figure 5B, right panel). Pitch was more weakly encoded in the ambiguous tones than the unambiguous tones: decoding in pure tones was significantly higher from 80-115ms (df = 27; t = 5.75, p < 0.001), and decoding in complex MF tones was significantly higher from 65-110ms (df = 27; t = 5.26, p < 0.001). However, the ability to decode pitch from these tones suggest that the auditory system settles on a pitch representation in the face of ambiguity, though weaker than that evoked by unambiguous tones.

Next, we assessed the spatial localization of pitch encoding. We used the same procedure described above, which was to estimate the decoding performance at each spatial source, for 50ms windows from −200 to 600ms. In pure tones (Figure 5A, left panel), pitch was encoded between 0-50ms in the left postcentral gyrus (df = 27; t = −2.06, p = 0.05), with significantly higher activity in sensorimotor areas than auditory (df = 27; t = 2.32, p = 0.03). Left Heschl’s gyrus significantly encoded pitch between 50-100ms (df = 27; t = 2.37, p = 0.03), and the right superior and middle temporal gyri showed significant encoding from 50-150ms (df = 27; t = 3.39, p = 0.005; df = 27; t = 2.89, p = 0.005). Heschl’s gyrus showed pitch encoding in the 250-300ms window (df = 27; t = 2.06, p = 0.05, with higher right hemisphere encoding during 300-350ms (df = 27; t = −2.33, p = 0.03). There was higher sensorimotor area encoding than auditory from 350-400ms (df = 27; t = 2.74, p = 0.01), with significant encoding in the left precentral and postcentral gyri (df = 27; t = 2.12, p = 0.04; df = 27; t = −2.45, p = 0.02).

In complex MF tones (Figure 5A, middle panel), pitch was encoded between −100--50ms before tone onset in left Heschl’s gyrus (df = 27; t = 2.08, p = 0.05). Right Heschl’s and middle temporal gyri encoded pitch between 50-100ms (df = 27; t = 2.68, p = 0.01; df = 27; t = 3.93, p = 0.0006), with bilateral superior temporal gyrus encoding (df = 27; t = 2.09, p = 0.05; df = 27; t = 4.84, p < 0.0001). From 100-150ms, pitch was encoded bilaterally in Heschl’s gyrus (df = 27; t = 3.46, p = 0.002) and in superior temporal gyrus (df = 27; t = 4.60, p = 0.0001). Similarly to pure tones, there was higher sensorimotor area encoding than auditory in later windows, from 350-450ms (df = 27; t = 3.32, p = 0.002; df = 27; t = 2.07, p = 0.05).

Ambiguous tone decoding showed a different pattern (Figure 5A, right panel), with pitch being encoded −50-0ms before tone onset in left Heschl’s and superior temporal gyri (df = 27; t = 2.44, p = 0.02; df = 27; t = 2.09, p = 0.05), and right middle temporal gyrus (df = 27; t = 2.83, p = 0.009). From 0-50ms, right Heschl’s gyrus encoded pitch (df = 27; t = 2.39, p = 0.02), pitch was not significantly encoded in any ROIs from 50-100ms, and right Heschl’s gyrus encoded pitch from 100-200ms (df = 27; t = 2.17, p = 0.04, df = 27; t = 2.59, p = 0.02). Later windows were dominated by sensorimotor areas, with significant pitch encoding in the right precentral gyrus from 250-300ms (df = 27; t = −2.31, p = 0.03), bilateral precentral and right postcentral gyri from 400-450ms (df = 27; t = −2.70, p = 0.01; df = 27; t = −2.57, p = 0.02, and right precentral gyrus from 500-550ms (df = 27; t = −2.63, p = 0.009). In sum, pitch encoding did not broadly alter its spatial distribution across tone types, but it was significantly attenuated in the ambiguous tones, mirroring the temporal decoding result. These findings suggest that pitch is robustly encoded in response to isolated F0, isolated harmonic structure, and ambiguous mixtures of these two cues leads to weaker pitch encoding.

### Role of context in pitch encoding

So far, the results focused on the encoding of two acoustic cues for pitch: fundamental frequency and inter-harmonic spacing. Next, we examine a third cue for pitch – expectation, given a tonal context.

We presented the pure and complex MF tones to our participants in a scale (high expectation) and random (low expectation) context. To assess the role of context for pitch encoding, we fit separate decoders on these two contexts and re-ran the temporal decoding procedure (Figure 5C, left and middle panels). In pure tones, the mean t-value for all clusters was smaller than 1.96 (uncorrected p < 0.05), showing the difference in decoding performance for scale vs. random context did not reach significance. In complex MF tones, decoding performance was significantly higher overall for the random context as opposed to the scale context from 85-115ms (df = 27; t = 3.52, p = 0.01).

Regarding pitch encoding outside the window of tone presentation, decoding performance separated by scale vs. random context in pure tones showed that pitch was significantly encoded before (−100 – −70ms; df = 27; t = 3.58, p = 0.04) and after (350 – 595ms; df = 27; t = 3.41, p = 0.0001) tone presentation when presented *in a scale context,* but not for tones presented randomly. Similarly, pitch was significantly encoded after tone presentation (415 – 445ms; df = 27; t = 3.08, p = 0.04; 555 – 575ms; df = 27; t = 3.74. p = 0.04) for complex MF tones presented in a scale context, but not when presented randomly. Thus, pitch encoding before and after tone presentation, was limited to the stimuli presented in a scale, and may reflect pitch prediction and maintenance during silent periods.

Next, we assessed the role of context for pitch encoding in the ambiguous tones, using the same decoding procedure. Pitch ambiguity has been shown to be resolved in context (Deutsch et al., 2008), so we created a high expectation condition: scales that end and begin on the same octave-ambiguous tone. When applying the analysis to the high expectation scale condition, we found that pitch can be decoded from −200 – −140ms (df = 27; t = 4.89, p = 0.004), 35-240ms (df = 27; t = 4.38, p = 0.0001), and 515-595ms (df = 27; t = 5.56, p = 0.0002); when applied to the low expectation random condition, pitch was encoded for a much shorter time period: from only 120 – 165ms (df = 27, t = 2.76, p = 0.003). We found that pitch could be decoded from the scale condition significantly better than the random condition from 55 - 110ms (df = 27; t = 2.93, p = 0.002; Figure 5C, right panel). Overall, these results indicate that pitch is decodable earlier when context is available as a cue to help derive the pitch percept of an ambiguous tone.

In sum, the outcome of our context analysis is that expectation only facilitates pitch encoding in the cases where acoustic cues are ambiguous. In the pure and complex MF tones, the acoustic cues are strong enough in pure/complex MF tones at their decoding peaks (85 vs. 95ms, respectively) to derive pitch regardless of context. The result showing higher decoding performance for complex MF tones in a random context is perplexing and requires further study for a clearer interpretation. In ambiguous tones, where the cues are weak or contradictory, the latency of pitch encoding is significantly modulated by context. High prediction scale contexts lead to pitch being immediately extracted from the sensory input (earliest peak at 75ms); whereas low prediction random contexts lead to pitch being extracted much later (earliest peak at 130ms).

### Pitch generalization

The data establish that representations of pitch unfold along a similar timecourse and localize to similar brain areas, when elicited by pure and complex MF tones, with weaker classification performance in ambiguous tones. Next, we test whether this pattern is consistent with the hypothesis that the neural coding scheme is shared across the two, or whether ‘fundamental frequency pitch’ is represented in a distinct neural pattern of activity from ‘inter-harmonic interval pitch.’

To evaluate this, we trained a ridge regression model to decode pitch within one tone type and evaluated how well that model was able to reconstruct pitch based on *responses to the other tone type.* Given our observation that the peak in decoding performance differed between pure and complex MF tones, it is possible that the neural processes that unfold for the two different tone types are offset in time. To find the optimal alignment, we performed a temporal generalization analysis (King and Dehaene, 2014), which allows us to find correspondences between neural representations that may be temporally offset.

This analysis allows us to test different hypotheses regarding the representation of pitch across tone types, and the relative latency with which pitch is derived in each case. These hypotheses are visually presented in schematic Figure 6A. Of most importance is whether (i) there is any generalization across tone types at all, and if so, (ii) with what relative latency difference.

**Figure 6.**
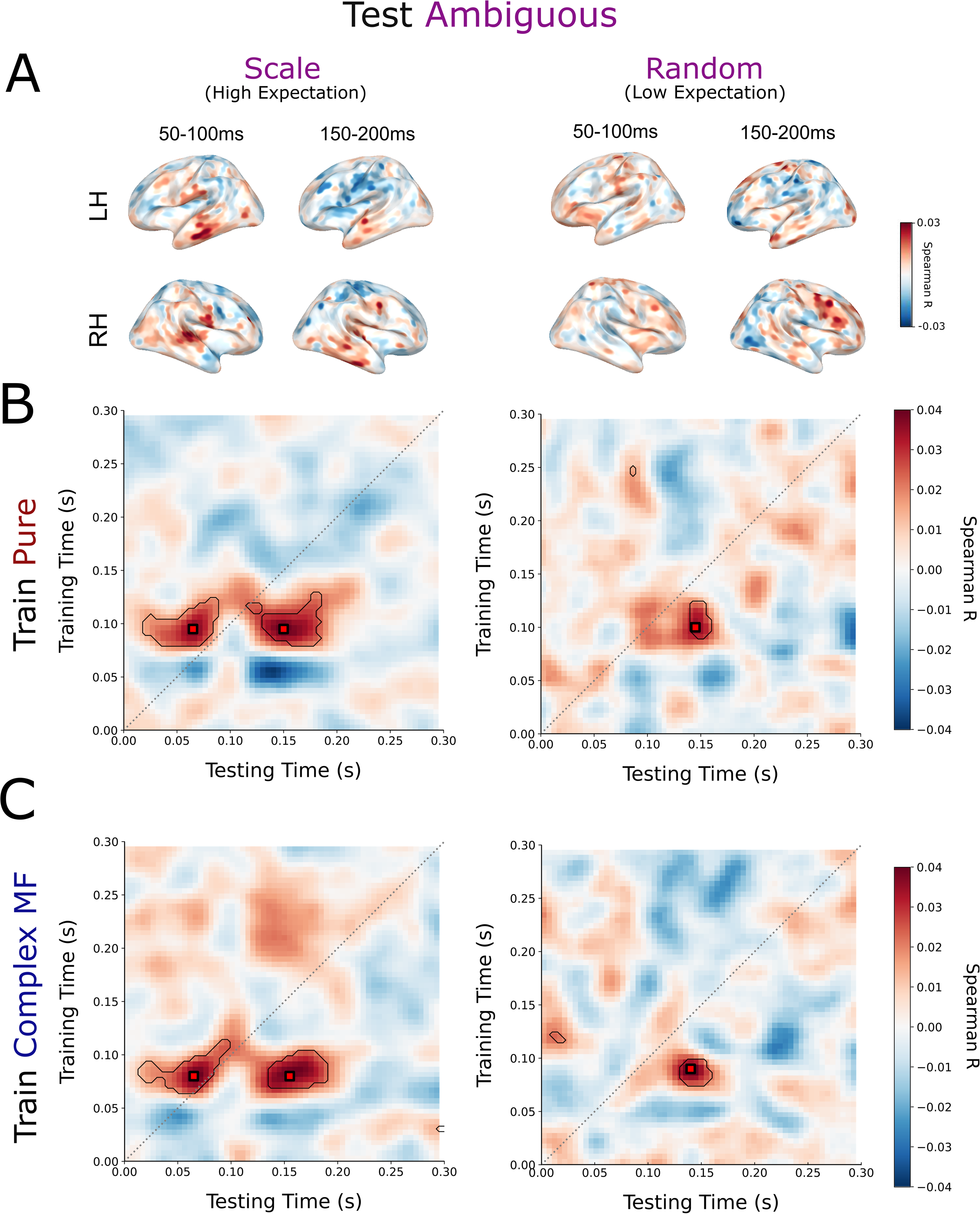
(A) Schematic of across-condition temporal generalization results under different neural hypotheses (top panel). If the neural pattern is distinct between conditions, at all latencies between conditions, we should observe no above-chance decoding at any train/test time pair. By contrast, if two conditions evoke shared neural representations, and that neural pattern is generated at the same latency across conditions, we will observe above-chance decoding performance along the diagonal, where training time is equal to testing time. If the same neural pattern is generated earlier in the test set as compared to the training set, we will observe above-chance decoding performance above the diagonal line. By contrast, if the same neural pattern is generated later, above-chance decoding performance will be below the diagonal. Finally, if the same neural pattern is generated transiently in the training set (e.g., pure tones), but maintained over time in the test set (e.g., we will observe above-chance decoding that begins at the diagonal and continues throughout test time. **Temporal generalization decoding for pitch within and across tone types (bottom panel). (B)** Decoding performance along the diagonal for pitch within pure tones and complex MF tones separately is plotted with standard error of the mean (*left*). Stimulus waveforms are depicted below each timecourse to denote tone duration. **(C)** Temporal generalization results for a model trained (y-axis) and tested (x-axis) on pure tones (*left*) and trained (y-axis) and tested (x-axis) on complex MF tones (*right*). Contours delineate t-value thresholds of 2 (gray dotted line) and 3 (black dotted line). **(D)** Decoding performance along a time-warped diagonal to adjust for latency differences in pitch decoding for a model trained on pure tones and tested on complex MF tones (red line) and vice versa (blue line; *left*). Stimulus waveforms are depicted below each timecourse to denote tone duration. **(E)** Temporal generalization results for a model trained (y-axis) on pure tones and tested (x-axis) on complex MF tones (*middle*) and trained (y-axis) on complex MF tones and tested (x-axis) on pure tones (*right*). Contours delineate t-value thresholds of 2 (gray dotted line) and 3 (black dotted line).

To establish an interpretable baseline for comparison, we first performed temporal generalization within each tone type (Figure 6B-C). As expected, there was robust decoding throughout the tone, with strongest decoding along the diagonal whereby the neural activity used to train the decoder best approximates the neural activity used to test the decoder.

Next, we applied the temporal generalization analysis *across* conditions. That is, we trained the decoder on pure tones and evaluated decoding performance on complex MF tones, and vice versa. This analysis revealed above-chance decoding that emerges earlier in complex MF tones (∼65-110ms, peaking at 80ms) than pure tones (∼75-140ms, peaking at 95ms) (df = 27; p = 0.0008, t = 2.99; Figure 4D-E). This suggests there is a *shared pitch representation* – the same neural pattern of activity encodes pitch across tone types – which is offset in time by ∼10ms. We use this optimal temporal alignment across tone type processing in subsequent analyses. Although the biological significance of this time offset is unclear, the complex MF tones acquire more pitch ‘evidence’ more quickly, as more neural ensembles are activated in response to multiple spectral components. There may be a number of underlying neural ensembles that give rise to this shared pattern.

Interestingly, although the initial pitch representation was shared across tone types during peak decoding time, we found that the later encoding of pitch, for example during the silent period between tones, did not generalize across conditions. This may suggest that while initial pitch encoding is shared between acoustic cues, the later representation is specific to the acoustic input that derived it.

### Combining acoustic and contextual pitch cues

Next, we tested whether the neural representation of pitch in the ambiguous tones is also shared with the pitch representation of the pure and complex MF tone stimuli, and whether this representation is affected by contextual expectation. Again, because the processing lag between stimulus types was not known *a priori*, we applied temporal generalization analyses, whereby we fit ridge regression decoding models on the responses to pure tones, and separately to complex MF tones, and then evaluated pitch decoding performance on the ambiguous tones. To evaluate the effect of context, we fit our decoders separately on the tones presented in a scale condition (providing high expectation contextual cues – “scale”) and those with random presentation order (low expectation contextual cues – “random”).

First, we tested if pitch is represented for ambiguous tones in the most predictive scale context. Our results revealed that the decoder, when trained on neural responses from 80-120ms to pure tones, significantly generalized to the representation of pitch in ambiguous tones both from 20-90ms (df = 27; t = 3.78, p = 0.0005) as well as from 125-180ms (df = 27; t = 3.51, p = 0.01). There were two distinct decoding peaks, with decoding of pure tones at 95ms generalizing most highly to ambiguous tones at 65ms (df = 27; t = 5.04, p < 0.001) and 150ms (df = 27; t = 4.45, p < 0.001), as shown in Figure 7B, left. A similar result was found for the complex MF tones: the pattern of neural activity from 60-105ms significantly generalized to ambiguous tones from both 15-90ms (df = 27; t = 3.86, p = 0.0006) and from 135-180ms (df = 27; t = 3.56, p = 0.0016). The two decoding peaks are similar but shifted forward in time across the training dimension by 15ms: decoding of complex MF tones at 80ms generalized most highly to ambiguous tones at 65ms (df = 27; t = 5.42, p < 0.001) and 155ms (df = 27; t = 3.93, p < 0.001), as shown in Figure 7C, left.

**Figure 7.**
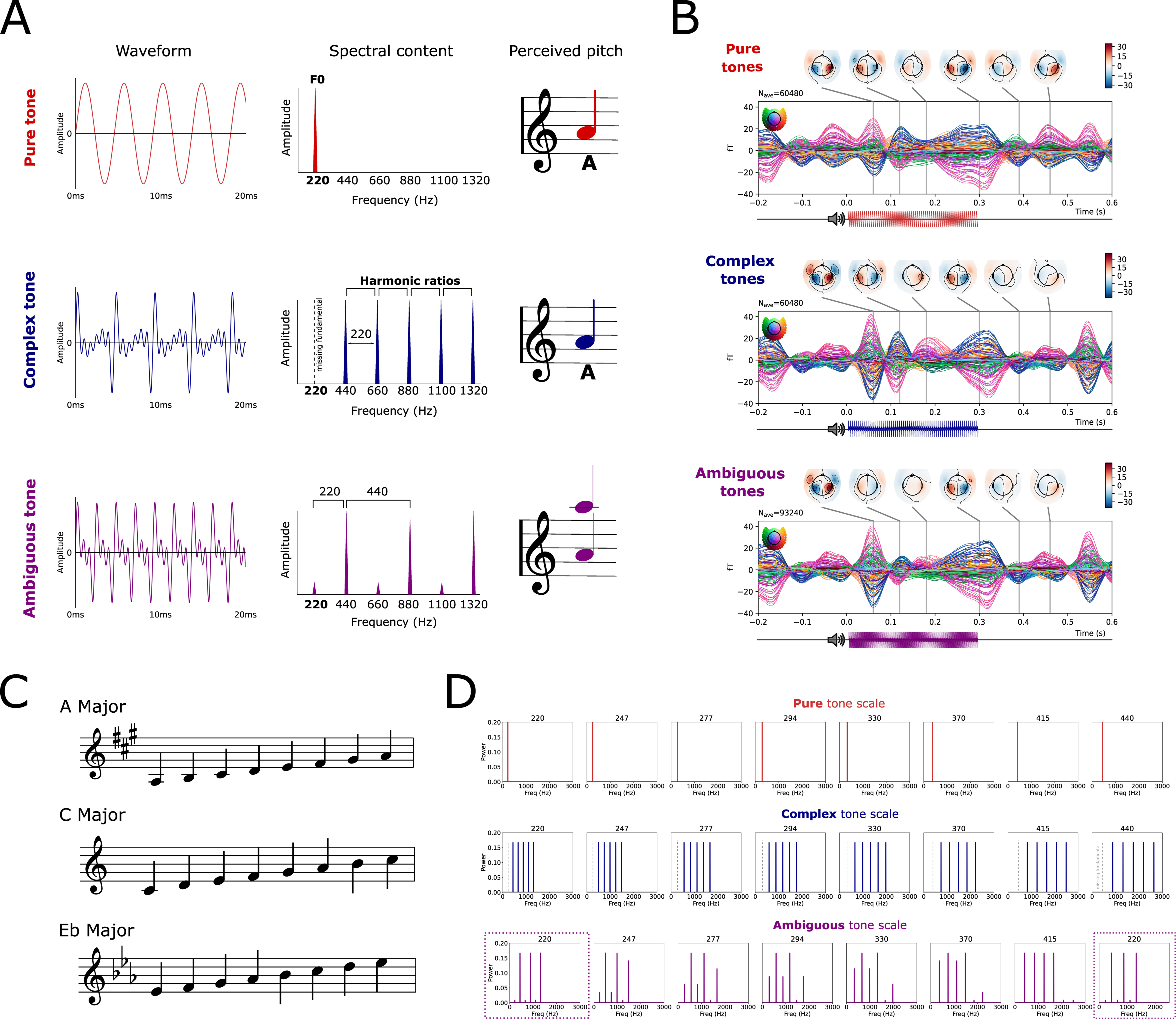
Temporal generalization for decoding pitch in ambiguous tones. **(A)** Source decoding for ambiguous tones in scale context (left) or random context (right) for the two windows in which pitch decoding from pure and complex tones are shown to be significant. **(B)** Temporal generalization results for decoding pitch from ambiguous tones presented in a *scale context* when trained on pure tones (left) or complex MF tones (right). Contours delineate a t-value threshold of 3 (thin black line). (**C)** Temporal generalization results for decoding pitch from ambiguous tones presented in a *random context* when trained on pure tones (left) or complex MF tones (right).

These results support two findings. First, the neural representation of pitch is not only shared between pure and complex MF tones but is also shared with ambiguous tones which contain a combination of pitch cues. Second, unlike the processing of pitch in unambiguous pure and complex MF tones, the pitch representation in ambiguous tones in context is maintained in neural activity during distinct early and later windows *throughout the duration of the tone*.

Finally, we sought to understand how the neural representation of pitch emerges over time when aiding context is removed for the ambiguous tone sequences. Interestingly, when applying our analysis to ambiguous tones presented in a random context, we found that the neural representation of pitch derived from pure tones during initial acoustic encoding at 90-120ms did not emerge in neural responses to ambiguous tones until 145-160ms (Figure 7B, right; df = 27; t = 3.54, p = 0.038). Decoding of pure tones at 100ms generalized with the highest performance to random ambiguous tones at 145ms (df = 27; t = 4.06, p < 0.0001). The neural representation of pitch derived from complex MF tones at 75-95ms generalized to 135-160ms (Figure 7C, right; df = 27; t = 4.03, p = 0.019). Decoding of pure tones at 90ms generalized with the highest performance to ambiguous tones at 140ms (df = 27; t = 5.35, p < 0.0001). In other words, *whereas in the scale presentation pitch is encoded from 20-90ms, when the same tones are presented in a random order, the same pitch cues are encoded, but not until 130-180ms*. This suggests that in the case where the acoustic cues are ambiguous, any context serves to generate an initial pitch representation, which is absent in the case where context is unreliable or considerably weaker.

We complement this generalization analysis by plotting pitch source decoding results within ambiguous tones for the two windows in which pitch generalized from pure and complex MF tones to ambiguous tones (50-100ms and 150-200ms), for each context (Figure 7A). This demonstrates a lack of any pitch representation in the early window (50-100ms) for randomly presented ambiguous tones, in contrast with representations of pitch in both the early and later window for ambiguous tones presented in a scale context.

## DISCUSSION

This study investigated how acoustic and contextual cues are cortically processed and integrated to facilitate pitch perception. We created stimuli that cued pitch via unambiguous fundamental frequency, unambiguous inter-harmonic intervals, and an ambiguous mixture, and we placed those signals in high and low expectation contexts. The time-resolved neural decoding approach we use provides a new analytic methodology to assess different possible representational schemes in the processing and representation of pitch.

We first address whether is pitch encoded differently for sounds with different spectral content (reflecting the input), or whether is there a shared pitch representation (reflecting the percept). Overall, we find that pitch is robustly encoded in neural responses to each tone *type* (consistent with the literature), and that the representation is shared between different acoustic and contextual cues. This unified pitch representation emerges at different latencies, however, depending on the acoustic cues available as well as the strength of the supporting context.

We then address when and where this shared representation is generated. The two acoustic cues for pitch appear to converge on the same representation in cortex, predominantly localizing to the right superior and middle temporal gyrus and surrounding auditory areas. Given the limitations of source localization accuracy with MEG, it is possible that the responses we observe in the middle temporal gyrus emerge from a bleeding over of activity from the auditory cortex. This is supported by the observation that the pattern of evoked responses is qualitatively similar across these ROIs (see Figure 2). This dominance of pitch representations in the right hemisphere is broadly in accordance with other work (Zatorre and Belin, 2001; Zatorre et al., 2002; Jamison et al., 2005; Hyde et al., 2008; Zatorre, 2022). Recent studies using lesion network mapping have confirmed right temporal and subcortical areas as crucial neural substrates for music processing, and more specifically, detecting fine pitch differences (Sihvonen et al., 2016; Sihvonen et al., 2024). This hemispheric asymmetry may be driven by differences in local *spectral* or *temporal interconnectivity patterns* in auditory cortex between the right and left hemisphere, respectively (Obleser et al., 2008; Cha et al., 2016). This bottom up, or, input-driven, specialization, however, has been shown to interplay with top-down mechanisms, such as perceptual categorization (Tierney et al., 2013), learning (Dehaene-Lambertz et al., 2005; Möttönen et al., 2006), and selective attention (Brodbeck and Simon, 2022). However, pitch is not completely absent from the left hemisphere, suggesting that there are pitch-encoding neural populations bilaterally (Allen et al., 2017; Zatorre, 2022). Given the simplicity of the auditory signals used to elicit pitch percepts in this study, low-level temporal information relating to pitch may still be encoded in left auditory regions, to be integrated with top-down information later in the processing stream. Interestingly, while we found that the distinction between pure and complex MF tones was right lateralized, the distinction between both unambiguous tone types and ambiguous tones was bilateral, if not slightly left lateralized. This distinction was also most prevalent in sensorimotor areas, which has not been shown before. The relevance of sensorimotor cortex in distinguishing between subtle differences in spectral information remains to be explored.

Although the data suggest that the neural representation of pitch recruits the same neural populations across pitch cues, we found that these populations peaked significantly earlier for complex MF tones (∼85ms) than pure tones (∼95ms). This could be attributed to the fact that the complex MF tones contain higher-frequency spectral content, with spectral centroids closer to the shortest-latency ∼1 kHz response of neurons in auditory areas (Roberts and Poeppel, 1996; Roberts et al., 1998). Although this difference is small, is supported by work showing that both F0 and pitch height, i.e. the spectral envelope, influence the latency of the pitch response (Ritter et al., 2007; Sauvé et al., 2021). Alternatively, the latency difference we observe could be due the fact that the higher frequencies provide a larger number of completed frequency cycles (i.e. more evidence) in order to derive a frequency estimate (Gage and Roberts, 2000), or to pitch being faster to compute based on harmonic interval cues than fundamental frequency based on the acoustic cues available from the periphery. The mechanistic origin of the latency difference we find merits further targeted investigation.

Lastly, we explored how the pitch representation is affected by contextual expectations. We found that when the acoustic cues for pitch are clear and unambiguous, there is little effect of context, in either the strength of pitch encoding or its latency. In fact, we found a slight increase in pitch encoding in the random context for complex MF tones, which may be explained under a predictive coding framework, whereby the higher prediction error generated by randomly presented tones leads to stronger pitch representations. For ambiguous signals, however, we found that context played a major role in adjusting the dynamics of pitch processing. Concretely, pitch was decodable with a peak around 80ms when presented in a predictable context and around 120ms when presented in a random context. The immediacy of pitch representation in predictable contexts suggests that the *expectation* of pitch is instantly integrated with — or used to directly bias the processing of — the impoverished acoustic cues. This result fits with recent work showing that context does not necessarily enhance the pitch representation itself, but rather primes the auditory system for expected stimuli (Graves and Oxenham, 2017). Thus, the pitch of ambiguous tones was more easily accessible earlier when presented in context, but a similarly strong pitch representation still emerged, albeit later, without context. Listening context, as opposed to the presence of a tonal hierarchy, may play a more direct role in *enhancing sensitivity* to pitch (Garcia et al., 2010). This result also supports the mechanism of auditory prediction signaling, in which target predictions are encoded in auditory cortex (Chouiter et al., 2015; Berlot et al., 2018). Whether predictions are propagated down from higher cortical areas or up from subcortical activity remains to be elucidated. In addition to showcasing the importance of expectation in pitch processing, the influence of context on acoustically identical sounds demonstrated here also highlights the dynamic flexibility of the processing system and the ability to integrate different cues, at different times, depending on the information available.

The findings we report show that different cues for pitch, including acoustic and contextual information, converge on a shared neural representation that is normalized for differences in spectral content and expectation. In naturalistic settings, pitch will likely be derived by combining information across multiple cues, some weak and some strong. Our results indicate that regardless of the specific cues used to generate a pitch percept, the representation converges on a common activity pattern in right-lateralized superior temporal gyrus. This finding supports the existence of a pitch-processing center (Patterson et al., 2002; Krumbholz et al., 2003; Bendor and Wang, 2005; Chait et al., 2005). However, this does not preclude the presence of other frequency- or timbre-based representations throughout auditory cortex – high decoding of pitch *within* (but not across) tone types specifically during later time windows suggests spectrally specific tonotopic representations downstream (Allen et al., 2022). Importantly also, we did not fix the spectral bandwidth across stimulus types, as our interest was in comparing orthogonal cues to pitch. Thus, in our stimuli, not only does the musical pitch get modulated up and down the scale, but the spectral content (harmonics) is also shifted within the same range (one octave). Thus, decoding performance may reflect this relative spectral shift in addition to pitch. Additionally, high decoding of tone type (pure vs. complex MF), suggests the existence of a timbre-sensitive region within both auditory and sensorimotor cortex (Bizley et al., 2009; Allen et al., 2017). Similar analyses could be applied to an exploration of absolute vs. relative pitch encoding and the role of pitch for different auditory domains, such as melodic and speech processing.

## Acknowledgments

The authors thank Ben Lang for help with data collection. We are very grateful to Pablo Ripollés and David Poeppel for their feedback on this manuscript. This project received funding from the Abu Dhabi Research Institute G1001 (awarded to AM); BRAIN Foundation A-0741551370 (LG); Whitehall Foundation 2024-08-043 (LG)

## Conflict of interest

The authors declare no competing financial interests.

## Notes

### Competing Interest Statement

The authors have declared no competing interest.

### Summary of Updates

Introduction updated to include recent work on pitch using similar stimuli and methods. Additional Figure 2 added showing raw MEG activity in response to tones separated by pitch and key in auditory and sensorimotor areas. Additional Figure 3 and accompanying analyses added showing the variability in strength and decoding timecourse across individuals. Expanded source decoding to 50ms windows shown in Figures 4, 5, and 7. Author (Isaac Krementsov) added. Supplemental files removed.

## References

Adachi Y, Shimogawara M, Higuchi M, Haruta Y, Ochiai M (2001) Reduction of non-periodic environmental magnetic noise in MEG measurement by continuously adjusted least squares method. IEEE Transactions on Applied Superconductivity 11:669–672.

Allen EJ, Burton PC, Olman CA, Oxenham AJ (2017) Representations of Pitch and Timbre Variation in Human Auditory Cortex. The Journal of Neuroscience 37:1284.

Allen EJ, Mesik J, Kay KN, Oxenham AJ (2022) Distinct Representations of Tonotopy and Pitch in Human Auditory Cortex. J Neurosci 42:416–434.

Barbour DL, Wang X (2003) Contrast tuning in auditory cortex. Science 299:1073–1075.

Barker D, Plack CJ, Hall DA (2011) Reexamining the Evidence for a Pitch-Sensitive Region: A Human fMRI Study Using Iterated Ripple Noise. Cerebral Cortex 22:745–753.

Bendor D, Wang X (2005) The neuronal representation of pitch in primate auditory cortex. Nature 436:1161–1165.

Berlot E, Formisano E, De Martino F (2018) Mapping Frequency-Specific Tone Predictions in the Human Auditory Cortex at High Spatial Resolution. J Neurosci 38:4934–4942.

Bizley JK, Walker KM, Silverman BW, King AJ, Schnupp JW (2009) Interdependent encoding of pitch, timbre, and spatial location in auditory cortex. J Neurosci 29:2064–2075.

Brattico E, Näätänen R, Tervaniemi M (2001) Context Effects on Pitch Perception in Musicians and Nonmusicians: Evidence from Event-Related-Potential Recordings. Music Perception 19:199–222.

Briley PM, Breakey C, Krumbholz K (2012) Evidence for Pitch Chroma Mapping in Human Auditory Cortex. Cerebral Cortex 23:2601–2610.

Brodbeck C, Simon JZ (2022) Cortical tracking of voice pitch in the presence of multiple speakers depends on selective attention. Front Neurosci 16:828546.

Brosch M, Schreiner CE (2000) Sequence sensitivity of neurons in cat primary auditory cortex. Cerebral Cortex 10:1155–1167.

Brosch M, Schulz A, Scheich H (1999) Processing of sound sequences in macaque auditory cortex: Response enhancement. Journal of Neurophysiology 82:1542–1559.

Butler BE, Trainor LJ (2012) Sequencing the Cortical Processing of Pitch-Evoking Stimuli using EEG Analysis and Source Estimation. Frontiers in Psychology 3.

Cha K, Zatorre RJ, Schönwiesner M (2016) Frequency Selectivity of Voxel-by-Voxel Functional Connectivity in Human Auditory Cortex. Cereb Cortex 26:211–224.

Chait M, Poeppel D, Simon JZ (2005) Neural Response Correlates of Detection of Monaurally and Binaurally Created Pitches in Humans. Cerebral Cortex 16:835–848.

Cheung C, Hamilton LS, Johnson K, Chang EF (2016) The auditory representation of speech sounds in human motor cortex. eLife 5:e12577.

Chouiter L, Tzovara A, Dieguez S, Annoni JM, Magezi D, De Lucia M, Spierer L (2015) Experience-based Auditory Predictions Modulate Brain Activity to Silence as do Real Sounds. J Cogn Neurosci 27:1968–1980.

Criscuolo A, Pando-Naude V, Bonetti L, Vuust P, Brattico E (2022) An ALE meta-analytic review of musical expertise. bioRxiv:2021.2003.2012.434473.

Dale AM, Liu AK, Fischl BR, Buckner RL, Belliveau JW, Lewine JD, Halgren E (2000) Dynamic statistical parametric mapping: combining fMRI and MEG for high-resolution imaging of cortical activity. Neuron 26:55–67.

de Cheveigné A (2005) Pitch Perception Models. In: Pitch: Neural Coding and Perception (Plack CJ, Fay RR, Oxenham AJ, Popper AN, eds), pp 169–233. New York, NY: Springer New York.

de Cheveigné A (2010) Pitch perception. In: Oxford Handbook of Auditory Science: Hearing (Plack CJ, ed): Oxford University Press.

Dehaene-Lambertz G, Pallier C, Serniclaes W, Sprenger-Charolles L, Jobert A, Dehaene S (2005) Neural correlates of switching from auditory to speech perception. Neuroimage 24:21–33.

Deutsch D (1992) Paradoxes of Musical Pitch. Scientific American 267:88–95.

Deutsch D, Dooley K, Henthorn T (2008) Pitch circularity from tones comprising full harmonic series. J Acoust Soc Am 124:589–597.

Englitz B, Akram S, Elhilali M, Shamma S (2024) Decoding contextual influences on auditory perception from primary auditory cortex. bioRxiv:2023.2012.2024.573229.

Gage NM, Roberts TP (2000) Temporal integration: reflections in the M100 of the auditory evoked field. Neuroreport 11:2723–2726.

Galbraith GC (1994) Two-channel brain-stem frequency-following responses to pure tone and missing fundamental stimuli. Electroencephalography and Clinical Neurophysiology/Evoked Potentials Section 92:321–330.

Gander PE, Kumar S, Sedley W, Nourski KV, Oya H, Kovach CK, Kawasaki H, Kikuchi Y, Patterson RD, Howard MA, Griffiths TD (2019) Direct electrophysiological mapping of human pitch-related processing in auditory cortex. NeuroImage 202:116076.

Garcia D, Hall DA, Plack CJ (2010) The effect of stimulus context on pitch representations in the human auditory cortex. NeuroImage 51:808–816.

Gordon CL, Cobb PR, Balasubramaniam R (2018) Recruitment of the motor system during music listening: An ALE meta-analysis of fMRI data. PLOS ONE 13:e0207213.

Gordon EM et al. (2023) A somato-cognitive action network alternates with effector regions in motor cortex. Nature 617:351–359.

Grahn JA, Rowe JB (2009) Feeling the beat: premotor and striatal interactions in musicians and nonmusicians during beat perception. J Neurosci 29:7540–7548.

Gramfort A, Luessi M, Larson E, Engemann DA, Strohmeier D, Brodbeck C, Goj R, Jas M, Brooks T, Parkkonen L, Hämäläinen M (2013) MEG and EEG data analysis with MNE-Python. Frontiers in Neuroscience 7.

Graves JE, Oxenham AJ (2017) Familiar Tonal Context Improves Accuracy of Pitch Interval Perception. Frontiers in Psychology 8.

Greenberg S (1980) WPP, No. 52: Temporal Neural Coding of Pitch and Vowel Quality. In.

Griffiths TD, Hall DA (2012) Mapping Pitch Representation in Neural Ensembles with fMRI. The Journal of Neuroscience 32:13343.

Griffiths TD, Kumar S, Sedley W, Nourski KV, Kawasaki H, Oya H, Patterson RD, Brugge JF, Howard MA (2010) Direct recordings of pitch responses from human auditory cortex. Curr Biol 20:1128–1132.

Hall DA, Plack CJ (2008) Pitch Processing Sites in the Human Auditory Brain. Cerebral Cortex 19:576–585.

Hall DA, Plack CJ (2009) Pitch Processing Sites in the Human Auditory Brain. Cerebral Cortex 19:576–585.

Hyde KL, Peretz I, Zatorre RJ (2008) Evidence for the role of the right auditory cortex in fine pitch resolution. Neuropsychologia 46:632–639.

Jamison HL, Watkins KE, Bishop DVM, Matthews PM (2005) Hemispheric Specialization for Processing Auditory Nonspeech Stimuli. Cerebral Cortex 16:1266–1275.

Kim S-G, Overath T, Sedley W, Kumar S, Teki S, Kikuchi Y, Patterson R, Griffiths TD (2022) MEG correlates of temporal regularity relevant to pitch perception in human auditory cortex. NeuroImage 249:118879.

King JR, Dehaene S (2014) Characterizing the dynamics of mental representations: the temporal generalization method. Trends Cogn Sci 18:203–210.

Krumbholz K, Patterson RD, Seither-Preisler A, Lammertmann C, Lütkenhöner B (2003) Neuromagnetic evidence for a pitch processing center in Heschl’s gyrus. Cereb Cortex 13:765–772.

Krumhansl CL, Shepard RN (1979) Quantification of the hierarchy of tonal functions within a diatonic context. In, pp 579–594. US: American Psychological Association.

Krumhansl CL, Cuddy LL (2010) A theory of tonal hierarchies in music. In: Music perception., pp 51–87. New York, NY, US: Springer Science + Business Media.

Kumar S, Sedley W, Nourski KV, Kawasaki H, Oya H, Patterson RD, Howard MA, III, Friston KJ, Griffiths TD (2011a) Predictive Coding and Pitch Processing in the Auditory Cortex. Journal of Cognitive Neuroscience 23:3084–3094.

Kumar S, Sedley W, Nourski KV, Kawasaki H, Oya H, Patterson RD, Howard MA, 3rd, Friston KJ, Griffiths TD (2011b) Predictive coding and pitch processing in the auditory cortex. J Cogn Neurosci 23:3084–3094.

Langner G, Sams M, Heil P, Schulze H (1998) Frequency and periodicity are represented in orthogonal maps in the human auditory cortex: Evidence from magnetoencephalography. Journal of comparative physiology A, Sensory, neural, and behavioral physiology 181:665–676.

Licklider JCR (1954) “Periodicity” Pitch and “Place” Pitch. The Journal of the Acoustical Society of America 26:945–950.

Lima CF, Krishnan S, Scott SK (2016) Roles of Supplementary Motor Areas in Auditory Processing and Auditory Imagery. Trends in Neurosciences 39:527–542.

Loui P, Wu EH, Wessel DL, Knight RT (2009) A Generalized Mechanism for Perception of Pitch Patterns. The Journal of Neuroscience 29:454–459.

Margulis EH, Mlsna LM, Uppunda AK, Parrish TB, Wong PCM (2009) Selective neurophysiologic responses to music in instrumentalists with different listening biographies. Human Brain Mapping 30:267–275.

Moerel M, De Martino F, Santoro R, Yacoub E, Formisano E (2015) Representation of pitch chroma by multi-peak spectral tuning in human auditory cortex. Neuroimage 106:161–169.

Monahan PJ, de Souza K, Idsardi WJ (2008) Neuromagnetic Evidence for Early Auditory Restoration of Fundamental Pitch. PLOS ONE 3:e2900.

Möttönen R, Calvert GA, Jääskeläinen IP, Matthews PM, Thesen T, Tuomainen J, Sams M (2006) Perceiving identical sounds as speech or non-speech modulates activity in the left posterior superior temporal sulcus. Neuroimage 30:563–569.

Nastase S, Iacovella V, Hasson U (2014) Uncertainty in visual and auditory series is coded by modality-general and modality-specific neural systems. Hum Brain Mapp 35:1111–1128.

Norman-Haignere S, Kanwisher N, McDermott JH (2013) Cortical pitch regions in humans respond primarily to resolved harmonics and are located in specific tonotopic regions of anterior auditory cortex. Journal of Neuroscience 33:19451–19469.

Obleser J, Eisner F, Kotz SA (2008) Bilateral speech comprehension reflects differential sensitivity to spectral and temporal features. J Neurosci 28:8116–8123.

Oxenham AJ (2012) Pitch perception. J Neurosci 32:13335–13338.

Oxenham AJ, Wojtczak M (2010) Frequency selectivity and masking. The Oxford handbook of auditory science: Hearing:5–44.

Pantev C, Hoke M, Lütkenhöner B, Lehnertz K (1989) Tonotopic organization of the auditory cortex: pitch versus frequency representation. Science 246:486–488.

Pantev C, Roberts LE, Elbert T, Roβ B, Wienbruch C (1996) Tonotopic organization of the sources of human auditory steady-state responses. Hearing Research 101:62–74.

Patterson RD, Uppenkamp S, Johnsrude IS, Griffiths TD (2002) The processing of temporal pitch and melody information in auditory cortex. Neuron 36:767–776.

Pedregosa F, Varoquaux G, Gramfort A, Michel V, Thirion B, Grisel O, Blondel M, Prettenhofer P, Weiss R, Dubourg V, Vanderplas J, Passos A, Cournapeau D, Brucher M, Perrot M, Duchesnay E (2011) Scikit-learn: Machine Learning in Python. Journal of Machine Learning Research 12:2825–2830.

Penagos H, Melcher JR, Oxenham AJ (2004) A neural representation of pitch salience in nonprimary human auditory cortex revealed with functional magnetic resonance imaging. Journal of Neuroscience 24:6810–6815.

Plack C, Fay R, Oxenham A, Popper A (2005) Pitch: Neural Coding and Perception.

Ritter S, Dosch HG, Specht H-J, Schneider P, Rupp A (2007) Latency effect of the pitch response due to variations of frequency and spectral envelope. Clinical Neurophysiology 118:2276–2281.

Roberts TP, Poeppel D (1996) Latency of auditory evoked M100 as a function of tone frequency. Neuroreport 7:1138–1140.

Roberts TP, Ferrari P, Poeppel D (1998) Latency of evoked neuromagnetic M100 reflects perceptual and acoustic stimulus attributes. Neuroreport 9:3265–3269.

Sauvé SA, Cho A, Zendel BR (2021) Mapping Tonal Hierarchy in the Brain. Neuroscience 465:187–202.

Schouten JF (1938) The perception of subjective tones. Proceedings of the Koninklijke Nederlandse Akademie van Wetenschappen 41:1086–1093.

Scott DW (1992) Multivariate density estimation: theory, practice, and visualization: Wiley & Sons, New York.

Seebeck A (1841) Beobachtungen über einige Bedingungen der Entstehung von Tönen. Annalen der Physik 129:417–436.

Shepard RN (1964) Circularity in Judgments of Relative Pitch. The Journal of the Acoustical Society of America 36:2346–2353.

Shera CA, Guinan JJ, Jr., Oxenham AJ (2002) Revised estimates of human cochlear tuning from otoacoustic and behavioral measurements. Proc Natl Acad Sci U S A 99:3318–3323.

Shera CA, Guinan JJ, Jr., Oxenham AJ (2010) Otoacoustic estimation of cochlear tuning: validation in the chinchilla. J Assoc Res Otolaryngol 11:343–365.

Sihvonen AJ, Ripollés P, Leo V, Rodríguez-Fornells A, Soinila S, Särkämö T (2016) Neural Basis of Acquired Amusia and Its Recovery after Stroke. J Neurosci 36:8872–8881.

Sihvonen AJ, Ferguson MA, Chen V, Soinila S, Särkämö T, Joutsa J (2024) Focal Brain Lesions Causing Acquired Amusia Map to a Common Brain Network. J Neurosci 44.

Slana A, Repovš G, Fitch WT, Gingras B (2016) Harmonic context influences pitch class equivalence judgments through gestalt and congruency effects. Acta Psychol (Amst) 166:54–63.

Thaut MH, Trimarchi PD, Parsons LM (2014) Human brain basis of musical rhythm perception: common and distinct neural substrates for meter, tempo, and pattern. Brain Sci 4:428–452.

Tierney A, Dick F, Deutsch D, Sereno M (2013) Speech versus song: multiple pitch-sensitive areas revealed by a naturally occurring musical illusion. Cereb Cortex 23:249–254.

Wallmark Z, Iacoboni M, Deblieck C, Kendall RA (2018) Embodied Listening and Timbre: Perceptual, Acoustical, and Neural Correlates. Music Perception 35:332–363.

Wang X, Walker KM (2012) Neural mechanisms for the abstraction and use of pitch information in auditory cortex. J Neurosci 32:13339–13342.

Warren JD, Uppenkamp S, Patterson RD, Griffiths TD (2003) Separating pitch chroma and pitch height in the human brain. Proceedings of the National Academy of Sciences 100:10038–10042.

Wilson SJ, Lusher D, Wan CY, Dudgeon P, Reutens DC (2009) The neurocognitive components of pitch processing: insights from absolute pitch. Cereb Cortex 19:724–732.

Zatorre RJ (2022) Hemispheric asymmetries for music and speech: Spectrotemporal modulations and top-down influences. Frontiers in Neuroscience 16.

Zatorre RJ, Belin P (2001) Spectral and Temporal Processing in Human Auditory Cortex. Cerebral Cortex 11:946–953.

Zatorre RJ, Belin P, Penhune VB (2002) Structure and function of auditory cortex: music and speech. Trends Cogn Sci 6:37–46.

Zatorre RJ, Chen JL, Penhune VB (2007) When the brain plays music: auditory-motor interactions in music perception and production. Nat Rev Neurosci 8:547–558.

